# Large-scale generation of uniform sub-100 μm adipocyte spheroids in hydrogel microcapsules using a flow-focusing microfluidic device

**DOI:** 10.1101/2024.06.08.597376

**Authors:** Ruri Maekawa, Kazuki Hattori, Hiromi Kirisako, Yuichiro Iwamoto, Fumiko Kawasaki, Takeshi Yoneshiro, Juro Sakai, Sadao Ota

## Abstract

Adipocyte spheroids are a promising three-dimensional (3D) cell culture model for obesity research because they reproduce 3D adipose tissue structures and cell-cell interactions better than 2D cultures. However, current methods fail to produce uniformly sized, small adipocyte spheroids at large scales, significantly limiting their use in analysis such as large-scale drug screening. Here, we develop a scalable method that combines simple microfluidics with templated emulsification to generate small, uniformly sized adipocyte spheroids. By encapsulating preadipocytes in numerous hollow agarose microcapsules and incubating them for two days, we reproducibly produced more than 100,000 uniform spheroids with diameters of approximately 50 µm (CV: <13%); we then differentiated preadipocyte spheroids into adipocyte spheroids after an 8-day induction period. Our platform enhances large-scale 3D analysis using adipocyte spheroids for obesity research and can be adapted to generate various spheroid and organoid models, advancing biomedical research across diverse fields.

## Introduction

Spheroids are essential models that better replicate the physiology of three-dimensional (3D) structures and cell-cell interactions of *in vivo* tissues compared to two-dimensional (2D) cell cultures (Barbosa et al., 2022; Białkowska et al., 2020). Differentiated spheroids, such as hepatocyte, neural, and cardiac spheroids, facilitate drug discovery through drug screening and analyzing mechanisms of action (Beauchamp et al., 2020; Hurrell et al., 2020; Strong et al., 2023). Additionally, spheroids generated from patient-derived induced pluripotent stem (iPS) cells enable the study of pathophysiology and the assessment of drug efficacy and toxicity in a more disease-relevant context (Plummer et al., 2019; Rowe and Daley, 2019). Among these, adipocyte spheroids have been thought valuable phenotypic models to develop methods to prevent or treat obesity, as they represent adipose tissues more accurately with increased levels of adipocyte marker expressions and lipid accumulations compared to 2D cultures (Klingelhutz et al., 2018; Shen et al., 2021; Turner et al., 2015, 2017).

To perform large-scale analyses with adipocyte spheroids, scalable technologies for generating small (<100 µm), uniformly sized spheroids hold great potential. Mass production of spheroids is crucial because large-scale assays, such as drug screening, typically require many spheroids to test thousands to millions of compounds independently (Macarron et al., 2011; Wang and Jeon, 2022). Small spheroids (<100 µm) are important for reducing culture costs and improving cell viability by minimizing cell death at the spheroid core. Furthermore, consistent spheroid size reduces variability in measurement outcomes, enhancing the reliability of the analytical platform.

Currently, no technology exists to produce uniformly sized small adipocyte spheroids on a large scale. Well-based methods, such as using 96-well or 384-well plates, face challenges in producing large quantities of spheroids (Amurgis et al., 2024; Graham et al., 2019). Additionally, due to the wide bottom area of the wells, forming spheroids typically requires hundreds or more cells, resulting in spheroids with diameters exceeding a few hundred micrometers. Hanging drop culture methods can create uniformly sized adipocyte spheroids but are labor-intensive and unsuitable for large-scale production (Akama et al., 2017; Robledo et al., 2023). While microwell platforms have expanded the number of spheroids per plate to a few thousand, the number produced is still insufficient for high-throughput screening (HTS), and achieving size uniformity is challenging due to inconsistencies in cell densities per microwell (Kim et al., 2021; Oka et al., 2019). Alternatively, microfluidic technologies facilitate the high-throughput production of spheroids within emulsions or microcapsules (Chan et al., 2013; Liu et al., 2020; van Loo et al., 2023); however, no one has yet successfully cultured adipocyte spheroids using these methods.

Here, we develop a scalable method for producing small, uniformly sized adipose spheroids by combining simple microfluidics with templated emulsification to form preadipocyte spheroids in hollow agarose microcapsules and inducing their long-term differentiation over 8 days (Figure 1). In this method, we first encapsulate preadipocytes within alginate hydrogel droplets using a focused microfluidic device at high throughput (> 10,000 droplets/min) and induce gelation inside the droplets to form core gels. We then encapsulate the core inside agarose droplets with a templated emulsification method (Hatori et al., 2018) and form hollow agarose capsules after solidifying the agarose and dissolving the alginate. Inside the hollow capsule, cells spontaneously form preadipocyte spheroids approximately 50 µm in diameter (CV: <13%) within 2 days, and spheroids are formed in more than 80% of the microcapsules. Through the stepwise application of differentiation-inducing media, the preadipocyte spheroids differentiate into adipocyte spheroids, characterized by lipid droplet accumulation. This platform consistently allows us to produce more than 100,000 uniformly sized small adipocyte spheroids in a single experiment. This platform can extend its capabilities beyond adipocyte spheroids to produce various other spheroid and organoid models, owing to two crucial features of agarose microcapsules: high permeability for long-term culture in diverse media and the adjustable size through device design modifications.

**Figure 1.**
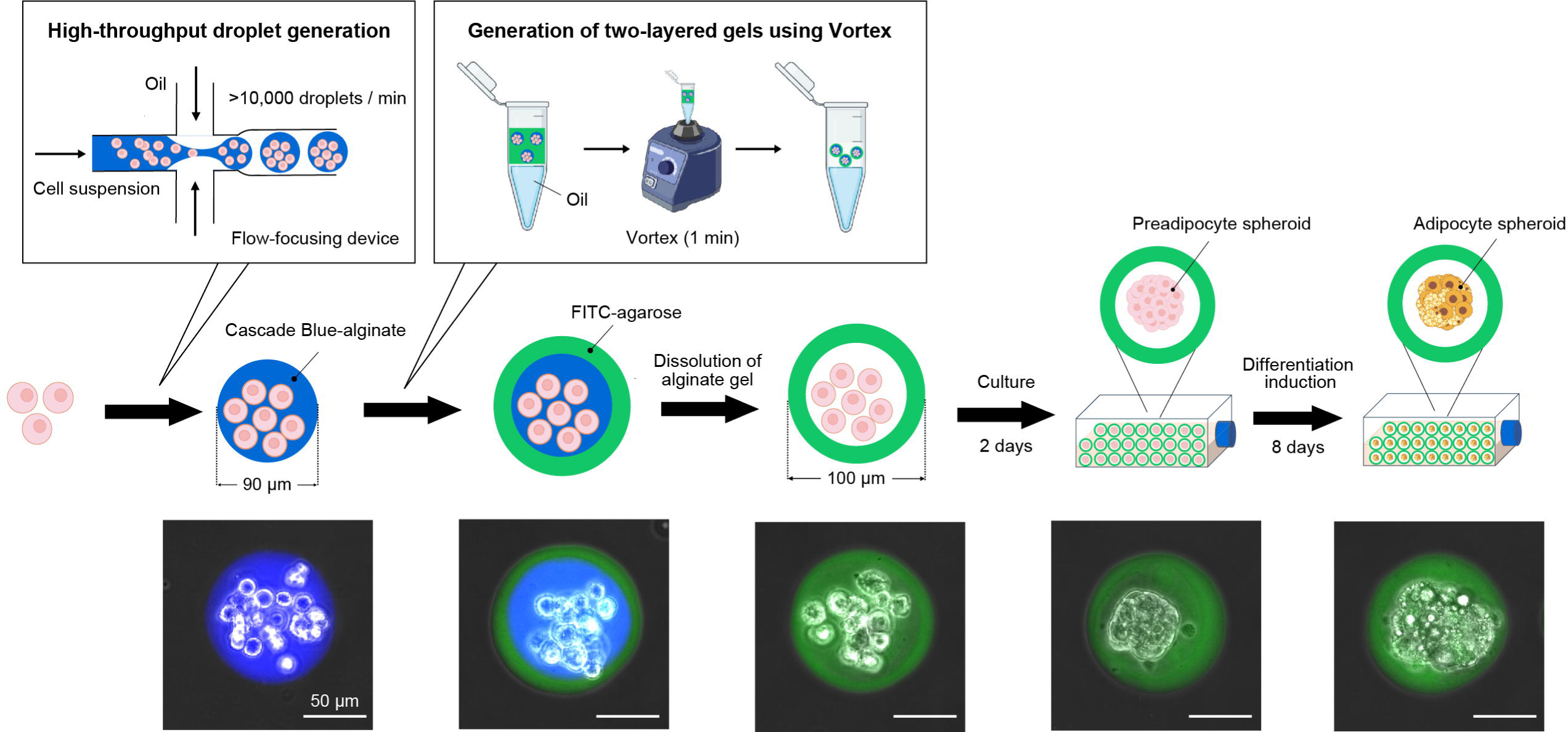
Large-scale generation of uniform sub-100 µm adipocyte spheroids in agarose microcapsules using droplet-based technologies. Preadipocytes are first encapsulated in alginate gel beads (Cascade Blue) using a flow-focusing microfluidic device fabricated using standard soft lithography. The alginate cores are then encapsulated inside agarose droplets (FITC) using a templated emulsification method to form the two-layered alginate-agarose gel structure. Washing the beads with calcium-free buffer, which dissolves only the alginate cores, results in the formation of hollow agarose microcapsules that enclose cells. After 2 days of culture, the cells spontaneously form preadipocyte spheroids. These spheroids then differentiate into adipocyte spheroids over the course of 8 days when exposed to differentiation-inducing media.

## Results

### Large-scale production of sub-100 **μ**m preadipocyte spheroids with uniform size and high cell viability in agarose microcapsules

To produce uniformly sized spheroids, we first aimed to generate uniformly sized hollow agarose microcapsules for culturing spheroids. We encapsulated an average of 15 preadipocytes within alginate core gels using a flow-focusing microfluidic device and subsequently produced hollow agarose microcapsules containing these cells through the templated emulsification method (Fig. 2A). To visualize the alginate core and agarose microcapsules, Cascade Blue was conjugated to the alginate and FITC to the agarose. Using a custom-made Python-based analysis platform, we calculated the area of each segmented core and microcapsule and determined their diameters, assuming a spherical shape. Analysis of 119 cores and 108 microcapsules indicated average diameters of 90.43 µm (CV = 1.72%) and 100.97 µm (CV = 3.07%), respectively. These results confirm the successful production of uniformly sized microcapsules that are suitable for spheroid culture.

**Figure 2.**
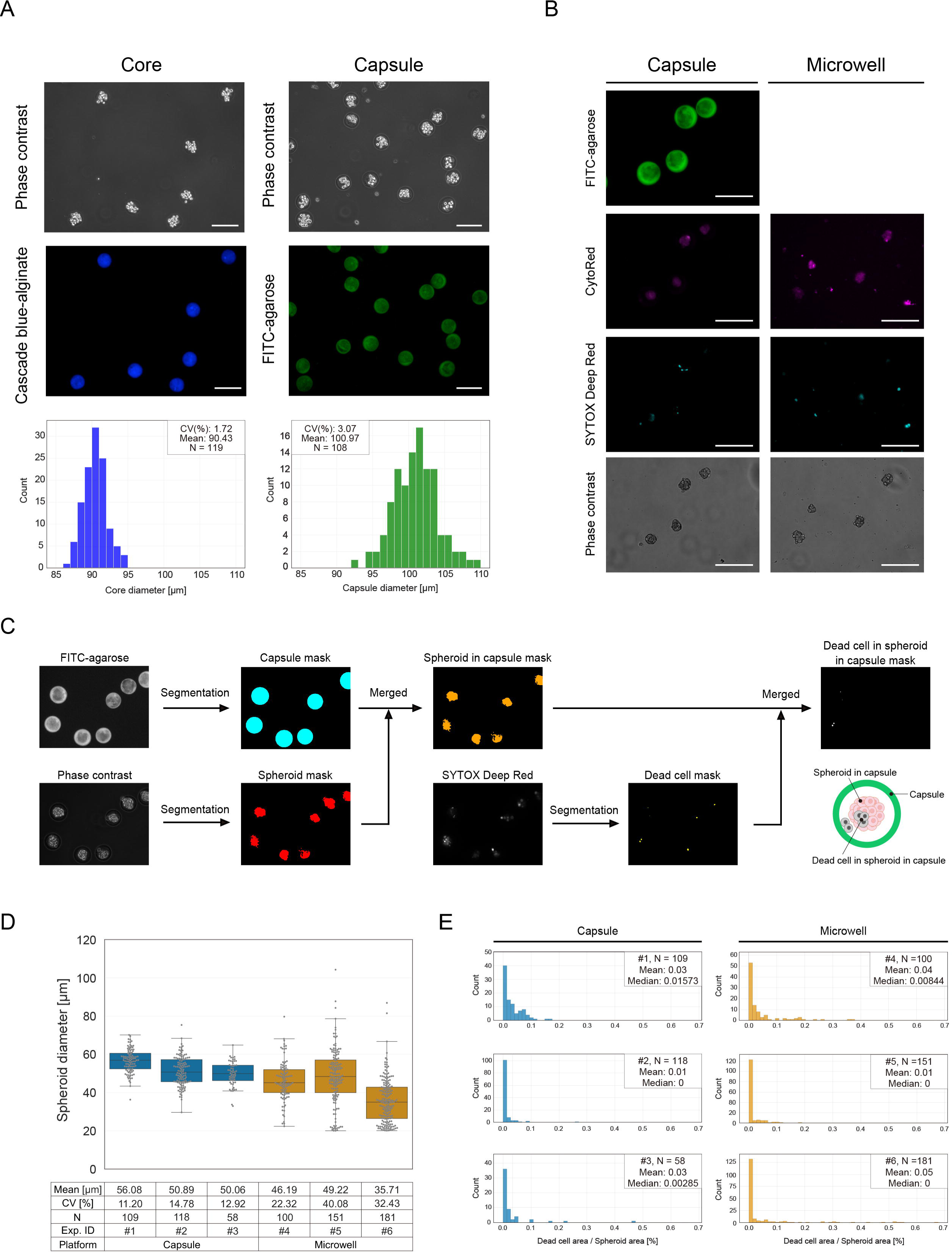
Large-scale production of uniformly sized small preadipocyte spheroids inside agarose microcapsules with high cell viability. (A) Representative fluorescence and phase contrast images of Cascade Blue-alginate cores and FITC-agarose microcapsules, with histograms of their diameters obtained by image analysis. Scale bar = 150 µm. (B) Representative fluorescence and phase contrast images of preadipocyte spheroids produced in the hydrogel-based microcapsules and microwells, 2 days after cell encapsulation or seeding. Green: microcapsule (FITC-agarose), magenta: live cells (CytoRed), cyan: dead cells (SYTOX Deep Red). Scale bar = 150 µm. (C) Workflow of the image analysis developed with Python 3. Refer to the method section for detailed information. (D, E) Quantification of the diameters of preadipocyte spheroids (D) and the levels of dead cells in spheroids (E) generated in microcapsules and microwells in three independent experiments. Diameters were calculated assuming a spherical shape and dead cell levels were obtained by dividing the dead cell area by the total spheroid area for each spheroid.

To confirm spheroid formation within the microcapsules, we conducted phase contrast and fluorescence imaging, using a microwell platform as a control method for producing spheroids. After two days of culturing cells in the microcapsules, we observed preadipocyte spheroids whose morphology was comparable to those produced in microwells (Fig. 2B). In this control experiment, we utilized Corning Elplasia plates in a 6-well format, each well containing about 3000 microwells with diameters of 500 µm and depths of 400 µm. We seeded a preadipocyte suspension at a concentration of 20 cells per microwell, which is slightly higher than the 15 cells per microcapsule used in the microcapsule method. This adjustment compensates for cells that do not settle at the bottom but remain suspended among the microwells. We then calculated the spheroid formation rate in microcapsules by dividing the number of microcapsules containing spheroids by the total number of microcapsules, achieving an average rate of 82.7% from three independent experiments. Furthermore, we replicated this workflow five times, consistently generating over 100,000 spheroids in each experiment, with a total processing time of approximately 2.5 hours per experiment.

Next, we evaluated the size uniformity and cell viability of preadipocyte spheroids cultured within the agarose microcapsules using spheroids produced by the microwell method as a control. Figure 2C illustrates the workflow of the image analysis developed with Python 3. We segmented the microcapsules and spheroids by binarizing FITC-agarose and phase-contrast images using Otsu’s method, respectively, and determined dead cell areas by binarizing SYTOX Deep Red images based on a manually defined threshold. We calculated the diameter of preadipocyte spheroids, assuming a spherical shape, and the dead cell ratio by dividing the dead cell area by the spheroid area for each spheroid. For the encapsulated samples, we omitted the spheroids outside of the microcapsules from the analysis. Analysis of three experimental replicates showed that the mean ± SD diameter of the spheroids generated in microcapsules was 52.3 ± 3.3 µm (CV: 13.0 ± 1.8%), while those in microwells was 43.7 ± 7.1 µm (CV: 31.6 ± 8.9%), suggesting that our microcapsule method can produce more uniformly sized spheroids (Fig. 2D). Additionally, dead cell ratios remained at 5% or less in all experiments, indicating high cell viability within the spheroids (Fig. 2E).

### Uniform adipocyte differentiation in agarose microcapsules

To demonstrate the compatibility of the agarose shell with differentiation induction, we induced adipogenic differentiation to preadipocyte spheroid over 8 days and confirmed that a high proportion of the cells in the preadipocyte spheroids differentiated into adipocytes, evidenced by the accumulation of lipid droplets. Figure 3A illustrates the differentiation protocol: spheroids were cultured in differentiation-inducing media for 8 days, with media changes every other day. Spheroids in microcapsules were centrifuged and resuspended in fresh differentiation media to replace all the media, whereas, in microwell plates, only half of the media was replaced to prevent spheroids from floating and subsequently merging.

**Figure 3.**
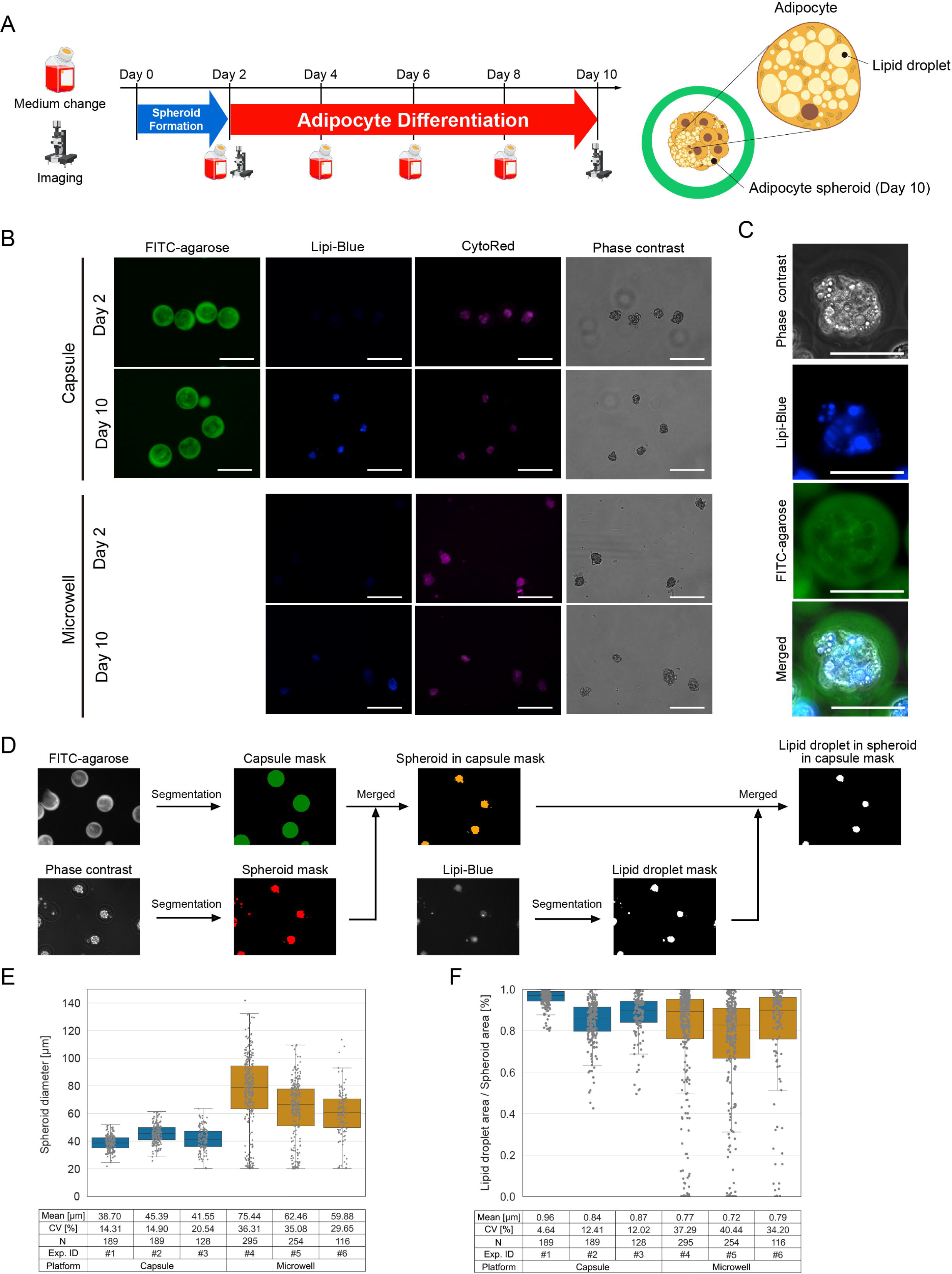
Uniform differentiation of adipose spheroids in agarose microcapsules. (A) Scheme of the adipocyte differentiation. Spheroids are formed in microcapsules or microwells from day 0 to day 2 in standard culture media. From day 2 to day 4, spheroids are cultured in differentiation induction media, and from day 4 to day 10 in differentiation maintenance media, with media changes every other day. Fluorescence and phase contrast imaging were performed on day 2 and day 10. (B, C) Representative fluorescence and phase contrast images of the adipocyte spheroids cultured in microcapsules and microwells on day 2 and day 10. Green: microcapsule (FITC-agarose), magenta: live cells (CytoRed), blue: lipid droplets (Lipi-Blue). Images in (B) are taken using a 20x objective lens, and those in (C) are taken using a 40x objective lens. Scale bar = 150 µm. (D) Workflow of the image analysis developed with Python 3. Refer to the method section for detailed information. (E, F) Quantification of the diameters of adipocyte spheroids (E) and the lipid accumulation levels (F) cultured in microcapsules and microwells. Diameters were obtained assuming a spherical shape and lipid accumulation levels were obtained by dividing the lipid droplet area by the total spheroid area for each spheroid.

To assess differentiation in adipocyte spheroids, we quantified lipid accumulation by measuring the area of lipid droplets within the spheroids and evaluated changes in spheroid size through the differentiation process. Lipid droplets were stained with a fluorescent dye (Lipi-Blue) and became clearly visible after 8 days of differentiation in both microcapsule and microwell samples (Fig. 3B and C). Using our Python-based image analysis platform, we quantified spheroid size and lipid accumulation levels. Following the method outlined in Figure 2C, we segmented the microcapsules and spheroids and identified regions of lipid droplets by binarizing the Lipi-Blue images using a manually defined threshold (Fig. 3D). We then calculated the diameter of the adipocyte spheroids, assuming a spherical shape, and the ratio of the lipid droplet area in each spheroid to assess differentiation.

First, we calculated the diameters of the spheroids. Spheroid diameters in microcapsules and microwells were 41.9 ± 3.4 µm and 65.9 ± 8.3 µm, respectively, calculated from three independent experiments. Additionally, we calculated the coefficient of variation (CV) in spheroid diameters within a single experiment and then calculated the mean ± SD across three independent experiments. The CV values were 16.6 ± 3.4% in microcapsules and 33.7 ± 3.5% in microwells. This confirms that the uniform size distribution of spheroids in microcapsules is maintained after differentiation, whereas spheroids in microwells exhibited higher size variation (Fig. 3E).

Next, we calculated the ratio of lipid droplet area per spheroid, similar to the analysis in Figure 3E. The ratio in microcapsules was 0.89 ± 0.06, while in microwell samples it was 0.76 ± 0.04. The CVs for these ratios were 9.7 ± 4.4% in microcapsules and 37.3 ± 3.1% in microwells. This measurement indicates that the microcapsule culture system achieved more uniform adipogenic differentiation compared to the microwell system (Fig. 3F).

## Discussion

Our method integrates simple microfluidics with templated emulsification to generate uniformly sized adipocyte spheroids within hollow agarose microcapsules. This technique consistently produces over 100,000 preadipocyte spheroids in 2.5 hours and induces differentiation into adipocyte spheroids over 8 days, demonstrating high throughput, reproducibility, and compatibility with long-term differentiation protocols.

Using agarose microcapsules approximately 100 µm in diameter, we produced small, uniformly sized spheroids with a diameter of approximately 50 µm. The confined core volume of approximately 0.5 nL ensures that most cells are in close proximity, promoting the formation of spheroids with uniform size. Traditional well-based platforms, such as the 96-well U-bottom plate, induce spheroid formation in a microliter-scale space, typically requiring hundreds or thousands of cells per well and resulting in spheroids larger than 200 µm in diameter (Ioannidou et al., 2022). Microwell platforms increase the number of spheroids per plate to several thousand by reducing the well volume to a sub-microliter scale, but spheroids tend to float and merge, increasing size variability (Kim et al., 2021; Razian et al., 2013). In contrast, our method isolates individual spheroids in microcapsules, preventing merging and subsequently reducing size variability.

Our small and uniformly sized spheroids offer several significant advantages to biomedical research. First, their smaller size minimizes cell death at the core by enhancing nutrient and oxygen supply (Wartenberg and Acker, 1995). Second, 3D imaging of numerous spheroids is feasible due to reduced measurement time per spheroid and reduced light scattering, facilitating high-throughput analysis and yielding more detailed images (Pampaloni et al., 2013). Lastly, our approach enables large-scale analysis of spheroids from rare cell types, such as patient-derived cells, since only a few dozen cells are needed to form a spheroid in microcapsules.

Using a simple flow-focusing device combined with templated emulsification, we achieved high-throughput and scalable spheroid production without requiring specialized skills. Unlike microwell platforms, which are limited to producing a few thousand spheroids per plate, our method enables the formation of over 100,000 spheroids in a single 2.5-hour experiment, with the potential for further scale-up through device parallelization. Additionally, culturing spheroids in microcapsules in a pooled format significantly increases the number of spheroids per unit volume of culture media, reducing the cost compared to well-based formats. This cost-effective and simple strategy for generating large numbers of spheroids enhances the feasibility of HTS.

The use of highly permeable agarose allows long-term culture of spheroids with high viability and efficient differentiation into adipocytes (Zarrintaj et al., 2018). The large pore size of agarose facilitates the supply of nutrients and reagents for differentiation within the microcapsules. Moreover, culturing spheroids in a suspended state allows easy media replacement through centrifugation and resuspension, whereas in microwell formats, only partial media changes are possible to avoid spheroid floating and merging. Although our current demonstration is limited to adipocyte spheroids, our platform is potentially advantageous for culture systems requiring multiple medium exchanges and prolonged differentiation induction, such as iPS cell-derived spheroids (Beauchamp et al., 2020; Castellanos-Montiel et al., 2023; Plummer et al., 2019).

We believe our platform has significant potential for HTS and 3D bioprinting applications. This potential is not limited to adipocyte spheroids but also includes liver, cardiac, and neural spheroids. HTS necessitates the production of large quantities of uniform and small spheroids for toxicity testing and drug efficacy screening. In 3D bioprinting, the uniformity and optimal size of spheroids are essential for constructing tissue structures with high spatial precision (Moor et al., 2018; Rawal et al., 2021). Our technology, which can mass-produce small, uniform spheroids, is adaptable for preparing materials for these applications, thereby enhancing both drug discovery and tissue engineering research.

## Experimental procedures

### Cell culture

We used immortalized preadipocytes derived from inguinal white adipose tissue (iWAT) for all experiments (Abe et al., 2018). iWAT preadipocytes were maintained in DMEM-high glucose (Wako, 045-30285) containing 10% fetal bovine serum (Sigma, F7524), 2 mM L-alanyl-L-glutamine (Wako, 016-21841), and Antibiotic-Antimycotic (100X) (Thermo Fisher Scientific, 15240096). This media is called culture media, hereafter. When the cells reached approximately 85% confluency, we passaged to a new 10 cm dish. Cells were cultured in a 5% CO_2_ atmosphere at 37°C. We confirmed the cells were mycoplasma-negative using a Cycleave PCR Mycoplasma Detection Kit (Takara Bio, CY232).

### Preparation of fluorescently labeled alginate

We first prepared a MES buffer (pH 5.5) by dissolving 100 mM MES hydrate (Sigma, M8250) and 300 mM NaCl in MilliQ water, adjusting the pH with 1 N NaOH. Next, we transferred 48 mL of this buffer to a 100 mL beaker. While stirring, we gradually added 50 mg of sodium alginate (Kimica Alginate IL-6) until the powder was fully dissolved. We then dissolved 100 μmol of N-ethyl-N’-(3-dimethylaminopropyl)carbodiimide hydrochloride (EDC, Merck, E7750) in 1 mL of MES buffer and added it to the beaker, stirring for 5 minutes. Subsequently, we added 1 mL of Cascade Blue solution (1 mg/mL in DMSO, Invitrogen, C687) and continued stirring in the dark overnight.

The following day, we aliquoted 10 mL of the sample from the beaker into 50 mL tubes, adding 40 mL of isopropanol (Wako, 166-04836) to each tube, followed by vortexing. The tubes were centrifuged at 3,000*g* for 5 minutes at room temperature. After discarding the supernatant, we evaporated the remaining isopropanol completely. The resulting pellets were then dissolved in 960 µL of MilliQ water per tube, combining the pellets from all five tubes into one, resulting in a total of 4.8 mL. This solution was aliquoted into twelve 2 mL tubes at 400 µL per tube, and each aliquot was mixed with 1.6 mL of isopropanol by vortexing. The tubes were then centrifuged at 10,000*g* for 5 minutes at room temperature. Following centrifugation, the supernatant was discarded, and the pellets were lyophilized.

To prepare the alginate solution for cell suspension, we first created a 2% alginate solution by dissolving sodium alginate in D-PBS(-). We also prepared an EDTA/Ca buffer containing 50 mM EDTA and 50 mM CaCl_2_ by diluting 1 M CaCl_2_ (Sigma, 21115-25L) and 0.5 M EDTA (pH 8.0, Thermo Fisher Scientific, AM9260G) in MilliQ water. We then combined the 2% alginate solution with the EDTA/Ca buffer in a 1:1 ratio, resulting in a final solution of 1% alginate, 25 mM EDTA, and 25 mM CaCl_2_.

### Preparation of fluorescently labeled agarose

We first dissolved Agarose Ultra-Low Gelling Temperature (Sigma, A5030-5G) in Dimethyl Sulfoxide (Super Dehydrated, Wako, 048-32811) at a concentration of 25 mg/mL, achieved through vortexing at room temperature. The vial containing this solution was sealed with a septum, and a needle was inserted to allow nitrogen gas to bubble through the solution. After bubbling, the vial was then placed on a ceramic hot stirrer (Asone, CHPS-170DF) set to 85°C for 4-5 hours to ensure complete evaporation of the DMSO.

For the FITC conjugation, we dissolved Fluorescein isothiocyanate isomer-I (Wako, F007) in DMSO at 100 mg/mL. An equal volume of agarose pellet was then added to the vial with agarose. The mixture was stirred on the ceramic hot stirrer set to 40°C using a stir bar for 1 hour. Following this, the agarose was lyophilized and reconstituted into a 1% solution with MilliQ water. This solution was then heated on the ceramic hot stirrer set to 55°C to ensure complete dissolution.

The FITC-agarose solution was transferred to 1.5 mL tubes, with each tube containing 1 mL of the solution. To remove unreacted reagents, these tubes were centrifuged at 20,000*g* for 3 minutes at 37°C. The supernatant was then transferred to new tubes and centrifuged 2-3 additional times until no sediment remained. The final FITC-agarose solution was aliquoted into 1.5 mL tubes and stored at 4°C.

For preparing the agarose solution for templated emulsification, we first prepared a 2% agarose solution by dissolving Agarose Ultra-Low Gelling Temperature in a buffer composed of 10 mM Tris (pH 8.0), 137 mM NaCl, 2.7 mM KCl, and 1.8 mM CaCl_2_. We then mixed the 1% FITC-agarose solution with the 2% agarose solution in a 3:97 ratio.

### Generation of spheroids in agarose shell

#### Cell preparation

After detaching the cells from 10 cm dishes and resuspending them in PBS, we added 20 µg/mL Deoxyribonuclease I (Wako, 047-26771) and gently agitated the suspension for 15 minutes at room temperature to minimize cell aggregation. The cells were then washed once with culture medium and centrifuged at 200*g* for 3 minutes to pellet them. After removing the supernatant, the cells were resuspended in the fluorescently labeled alginate solution to achieve a cell concentration of 5.6 × 10^7^ cells/mL, ensuring an average of 15 cells per droplet.

#### Microfluidic device setup

Using standard soft lithography, we fabricated flow-focusing devices with polydimethylsiloxane (PDMS, SILPOT184, DOW). The device has three inlets: one for the cell suspension, one for Automated Droplet Generation Oil for EvaGreen (Bio-Rad, 1864112) containing 0.05% acetic acid (Wako, 017-00256), and one for HFE7200 (3M) containing 20% 1H,1H,2H,2H-perfluoro-1-octanol (PFO, Wako, 324-90642). In part of this work, HFE7200 was replaced with HFE7500.

#### Generation of core alginate gels with a flow-focusing microfluidic device

The cell suspension was introduced into the microfluidic channels through PEEK tubing using FLPG Plus (Fluigent) and LineUp Flow EZ (Fluigent, LU-FEZ-2000) at 325 mbar. Droplet Generation Oil and HFE7200 or HFE7500 containing 20% PFO were introduced by a syringe pump (Harvard Apparatus, PUMP 11 ELITE) at flow rates of 35 µL/min and 25 µL/min, respectively. In this device, the alginate/EDTA/Calcium mixture solidifies immediately upon exposure to the acidic oil phase due to the release of calcium from EDTA, which occurs in response to the lowered pH and results in alginate crosslinking.

To minimize cell toxicity due to low pH, we broke the emulsions in flow by adding PFO-containing oil and collected the cell-containing alginate gels in a tube filled with culture media. Additionally, we changed the collection tubes every 5 minutes and accumulated the gel suspension in a new 15 mL tube (Corning, 352097) that contained 5 mL culture media to minimize the time of contact with acetic acid-containing oil.

#### Encapsulating core alginate gels in agarose droplets by templated emulsification

After centrifuging at 200*g* for 3 minutes and removing the supernatant, we added an equal volume of previously prepared fluorescently labeled agarose to the pellet.

Subsequently, Automated Droplet Generation Oil for EvaGreen, in a volume five times that of the alginate/agarose mixture, was added, and the tube was vortexed at maximum speed for 1 minute to achieve emulsification. The 15 mL tubes were then incubated on ice for 10 minutes to solidify the agarose gel, resulting in the formation of alginate-agarose two-layered gels with cells encapsulated within the alginate core. To break the emulsions, we combined them with HFE7200 containing 20% PFO and collected the gels in D-PBS(-) (Wako, 045-29795).

#### Create hollow agarose microcapsules and induce spheroid formation

To dissolve the alginate cores, we washed the two-layered gels four times with 10 mL of D-PBS(-) and once with 5 mL of culture media, with centrifugation at 200*g* for 3 minutes after each wash. We finally resuspended gels with DMEM-high glucose containing 20% fetal bovine serum, 4 mM L-alanyl-L-glutamine, and Antibiotic-Antimycotic (100X). and cultured them in 48-well plates (Corning, 353078) for two days.

### Generation of spheroids in Microwell

To generate spheroids in microwells, we used the Elplasia 6-well plates (Corning, 4440), which contain approximately 3000 microwells with diameters of 500 µm and depths of 400 µm. To remove air bubbles from the microwells before seeding the cells, 1.5 mL of culture media was added to each well, and the plate was centrifuged at 500*g* for 1 minute. This process was repeated twice to ensure complete bubble removal.

The cell suspension was adjusted so that each microwell contained approximately 20 cells. Each well was then filled with 2 mL of culture medium. The cell suspension was thoroughly mixed and distributed into the wells. Cells were cultured in a 5% CO_2_ atmosphere at 37°C for two days.

### Adipocyte differentiation

The induction of beige adipocytes was performed for both preadipocyte spheroids cultured in microcapsules and microwells. Two days after encapsulation in microcapsules or seeding in microwells (day 2), the culture media was replaced with differentiation induction media. This media consisted of culture media supplemented with 0.125 mM indomethacin (Sigma, I-7378), 2 μg/mL dexamethasone (Sigma, D-1756), 0.5 mM 3-isobutyl-1-methylxanthine (Sigma, I-5879), 0.5 μM rosiglitazone (Wako, 184-02651), 5 μg/mL insulin (Wako, 090-06481), and 1 μM T3 (Wako, 038-25541). On days 4, 6, and 8, the media was replaced with maintenance media, which contained only 0.5 μM rosiglitazone, 5 μg/mL insulin, and 1 μM T3.

For the microcapsule samples, we transferred the microcapsules to a 15 mL tube and centrifuged them at 200*g* for 3 minutes to remove the media. The microcapsules were then resuspended in fresh media. For the microwell samples, 1 mL of the 2 mL media was carefully removed from near the liquid level and replaced with 1 mL of fresh media. This process was repeated three times to ensure most of the media in the well was replaced with fresh medium.

### Fluorescence imaging

We performed imaging using the EVOS FL Auto Imaging System (Thermo Fisher Scientific, C10228) on day 2 for analyzing preadipocyte spheroids and on day 10 for mature adipocyte spheroids. Spheroids were collected in a 1.5 mL tube and centrifuged at 200g for 3 minutes. After removing the supernatant, the pellet was resuspended in 500 μL of DMEM high-glucose containing 0.2 μM Lipi-Blue (Dojindo Laboratories, LD01) and 0.1 μM SYTOX Deep Red Nucleic Acid Stain (Thermo Fisher Scientific, S11381). The samples were incubated in a 37°C, 5% CO_2_ atmosphere for 30 minutes for staining, followed by washing with FluoroBrite DMEM (Thermo Fisher Scientific, A1896701). The samples were then applied to the Countess Cell Counting Chamber Slides (Thermo Fisher Scientific, C10228) for imaging. For staining spheroids cultured in microwell plates, we pre-wetted the insides of micropipette tips and 1.5 mL microtubes with Cellotion (Nippon Zenyaku Kogyo, 13929) to prevent nonspecific absorption of the spheroids to the plastic surfaces. We performed imaging under consistent conditions for light intensity, exposure time, and gain within similar experiments. The recorded images are TIFF files with 12-bit grayscale resolution and a pixel size of 2048 x 1536.

### Image analysis

We developed the image analysis pipeline using SciPy and scikit-image in Python 3.10 (Virtanen et al., 2020; Walt et al., 2014).

#### Preprocessing

These images consist of five channels: Lipi-Blue for lipid droplets, FITC-agarose for microcapsules, CytoRed for live cells, SYTOX Deep Red for dead cells, and phase contrast. To reduce noise, we applied a Median filter and a Gaussian filter with a sigma value of 1 across all channels. We then applied erosion and dilation processes using a 5×5 square kernel to remove small artifacts and connect disjointed parts of objects.

#### Identification and analysis of core gels, microcapsules, and spheroids

To segment each alginate core gel, agarose microcapsule, and spheroid, we first binarized the images using Otsu’s thresholding method on the Cascade Blue-alginate, FITC-agarose, and phase contrast images, respectively. After binarization, we applied a hole-filling algorithm to ensure that the segmented regions were solid and removed objects touching the image border to exclude partial objects from the analysis. Individual gels, microcapsules, and spheroids were then labeled by applying the connected-component labeling method. Microcapsules were selected based on their circularity, calculated using the formula (4 * π * Area) / (Perimeter^2^), with a threshold value of 0.8.

To calculate the diameter of core gels, microcapsules, and spheroids, we extracted the area of each object using the regionprops function from the scikit-image package and calculated the diameter assuming a spherical shape using the formula: diameter = sqrt(Area / π) * 2.

For the encapsulated samples, we selected only spheroids within microcapsules by merging the segmented areas of spheroids and microcapsules to exclude spheroids cultured outside of the microcapsules from the analysis.

We excluded core gels and microcapsules with diameters less than 70 µm from the analysis in Figure 2A as they did not contain cells. We also excluded spheroids with diameters outside the range of 20 to 300 µm from the analysis in Figures 2D, 2E, 3E, and 3F to omit small cell clusters and artifacts.

#### Quantification of the spheroid formation rate

We calculated the spheroid formation rate by dividing the number of microcapsules containing spheroids by the total number of microcapsules. We segmented spheroids using CytoRed images using the same method used for phase contrast images. We considered only objects with a diameter of 30 µm or more as spheroids. Microcapsules were identified using FITC-agarose images as described above.

#### Identification and analysis of lipid droplets and dead cells

We segmented lipid droplets and dead cells by binarizing the Lipi-Blue and SYTOX Deep Red images using manually defined signal thresholds of 900 and 2000, respectively. We consistently applied these threshold values to all samples. We identified the area of lipid droplets in spheroids by merging the segmented areas of lipid droplets and spheroids.

We estimated lipid accumulation levels by dividing the lipid droplet area by the spheroid area for each spheroid. For the encapsulated samples, we selected only spheroids within microcapsules, as described above. We determined cell death levels similarly, replacing the lipid droplet area with the cell death area.

## Acknowledgment

We thank all the members of the networked biophotonics and microfluidics group. This work was supported by JST CREST grant number JPMJCR19H1 (to S.O.), JST GteX grant number JPMJGX23B1 (to S.O.), AMED-CREST grant number JP20gm1310007 (to T.Y. and J.S.), JSPS KAKENHI grant numbers JP21H04636 (to S.O.), and JP22K12797 (to K.H.), and AMED PRIME grant number JP22gm6710008 (to K.H.). This work was also supported by The Uehara Memorial Foundation (to S.O. and K.H.), UTEC-UTokyo FSI Research Grant Program (to S.O.), Takeda Science Foundation (to S.O. and K.H.), The Cell Science Research Foundation (to K.H.), Japan Society for the Study of Obesity (to K.H.), and Institute for Fermentation (to K.H.). A part of Figure 1 was created with BioRender.com.

## Author Contributions

Conceptualization, K.H. and S.O.; Methodology, K.H. and S.O.; Software, Y.I., K.H.; Validation, R.M. and H.K.; Formal Analysis, K.H., Y.I., and R.M; Investigation, R.M. H.K., and K.H.; Resources, K.H., S.O., F. K., T.Y., and J.S.; Data Curation, R.M and K.H.; Writing – Original Draft, R.M., K.H., and S.O.; Writing –Review & Editing, K.H., and S.O.; Visualization, R.M. and K.H.; Supervision, S.O. and K.H.; Project Administration, K.H. and S.O.; Funding Acquisition, S.O. and K.H.

